# Improving Protein Subcellular Localization Prediction with Structural Prediction & Graph Neural Networks

**DOI:** 10.1101/2022.11.29.518403

**Authors:** Geoffroy Dubourg-Felonneau, Arash Abbasi, Eyal Akiva, Lawrence Lee

## Abstract

The majority of biological functions are carried out by proteins. Proteins perform their roles only upon arrival to their target location in the cell, hence elucidating protein subcellular localization is essential for better understanding their function. The exponential growth in genomic information and the high cost of experimental validation of protein localization call for the development of predictive methods. We present a method that improves subcellular localization prediction for proteins based on their sequence by leveraging structure prediction and Graph Neural Networks. We demonstrate how Language Models, trained on protein sequences, and Graph Neural Networks, trained on protein’s 3D structures, are both efficient approaches for this task. They both learn meaningful, yet different representations of proteins; hence, ensembling them outperforms the reigning state of the art method. Our architecture improves the localization prediction performance while being lighter and more cost-effective.

## 1 Introduction

The elucidation of protein function is a fundamental challenge in biology. A significant aspect of protein function is the determination of its cellular localization. Many processes, such as disease mechanisms, drug performance, regulation of metabolic processes, and signaling cascades depend on the protein’s localization, hence significant experimental efforts are dedicated to this end. Nevertheless, the exponential growth in the availability of proteomics information from newly sequenced organisms and metagenomes creates a huge gap in the experimental elucidation of protein subcellular localization for gene and genome annotation. This creates a need to *predict* protein localization using sequence data.

Frequently, the location of a protein in the cell is determined by a “localization signal” – a short segment of the protein that is recognized by receptors located in the target cellular compartment. For example, proteins that include a *nuclear localization signal* (NLS) are recognized by nuclear receptors and, once synthesized, will be shuttled into the nucleus. Similarly, proteins that include a *signal peptide* will be exported to the extracellular space. Many computational methods are thus based on the identification of short motifs in the protein sequence that determines its cellular localization. Still, many proteins are shuttled to their target compartments using chaperones or other mechanisms, leading to an inaccurate prediction by motif-based approaches. More recent predictive models utilize the entire sequence of a protein to predict its localization using protein sequence language models such as Evolutionary Scale Modeling (ESM) [1]. The prediction power of these methods was shown to be higher than motif-based approaches [2]. The recent advance in protein structure prediction [3][4] enables another avenue of development: utilizing protein structure for the prediction of protein localization.

In this paper, we propose a Graph Neural Network (GNN) that utilizes protein structure information for protein subcellular localization prediction. We ensemble this model with a transformer-based language model that takes the protein sequence into account. We demonstrate that this combined method yields higher prediction performances than previous methods.

## 2 Background work

### 2.1 Language Model Representation

Meta AI’s ESM [1] is an unsupervised transformer protein language model. In this paper, we employ ESM-1b, in which the model is trained to predict amino acids from the surrounding sequence context, as the core model to extract abstract features from the protein sequences. Although a competing model – Hugging Face’s ProtT5XL [5][6] – is available, we focused on ESM-1b for its better ability to scale ^1^.

### 2.2 Subcellular localization

Alongside their cellular localization dataset, the authors of Deeploc 2 [2] published a successful Language Model (LM)-based classifier. It feeds the ESM-1b representation to a multi-class classifier involving an attention-pooling layer and a Multi-Label Perceptron (MLP). Its performance is summarized in Table 1.

**Table 1:** Summary of the classification metrics (micro average) on the HPA test set for different approaches. (1) GVP alone as a classifier (Figure 1.b), (2) LM baseline model (3 layers CNN), (3) Architecture proposed by Deeploc 2, (4) Our dense-residual ensembling (Figure 1), (5) A supplementary (non-differentiable) ensemble by the mean of 2. and 4. slightly outperforms 4. alone.

| Model | Precision | Recall | F1-score |
| --- | --- | --- | --- |
| 1. GVP alone | 0.34 | 0.69 | 0.46 |
| 2. LM baseline | 0.67 | 0.47 | 0.55 |
| 3. LM attention pooling (Deeploc 2) | - | - | 0.57 |
| 4. Proposed architecture (Figure 1) | 0.50 | 0.72 | 0.59 |
| 5. Ensemble of 2. and 4. | 0.59 | 0.60 | **0.60** |

## 3 Method

In recent years, the use of Language Models for protein representation has proven to be surprisingly successful for downstream prediction models; accordingly, their use has become increasingly prevalent in the field [1] [7] [5] [8]. Often, LMs are trained in a self-supervised fashion on hundreds of millions of proteins before being fine-tuned or used as backbone models for downstream tasks [9]. In parallel, the proteomics community underwent a major breakthrough with the inventions of very successful in-silico protein folders [3] [4]. The goal of our research is to combine those two new sources of information to outperform models that only utilize one such source.

### 3.1 Data

In this paper, we are benchmarking our method on the subcellular localization task made possible by the Deeploc 2 [2] dataset. This dataset is comprised of a train/validation set of 24674 protein sequences mapped to 10 localization classes (*e.g*. Cytoplasm, Nucleus, etc.), and a test set from a different source – the Human Protein Atlas (HPA)[10] – of 1,532 proteins associated with 8 localization classes ^2^, which we are using for testing.

For each protein entry, we retrieved the AlphaFold model from the AlphaFold Protein Structure Database [11] in the form of an atom coordinates file (PDB format). For each structure, we generated a graph representation based on the amino-acid adjacency matrix, inspired by the LM-GVP method [12]. The dihedral angle between amino acids is used as a node feature in the model to capture the geometrical relationships between the adjacent amino acids. We introduced two novel node attributes to the LM-GVP graph: (1) **positional encoding (PE)**: Adjacency matrices do not encode protein’s primary sequence, therefore (a) will have low chances of recognizing primary-sequence motifs, and (b) will be less sensitive to the distinction between short and long-range amino acid interactions, known to have different contributions to protein folding, structure and function [13], and (2) **AlphaFold’s Local Distance Difference Test (pLDDT)**: pLDDT is a metric of AlphaFold’s confidence for atom coordinate assignment. It has been observed that regions of the protein structure that have a higher pLDDT are generally associated to “less rigid” parts of the structure [14]. We hypothesized that pLDDT increases the predictive power for many downstream tasks, such as cellular localization.

### 3.2 Baseline

In order to compare the performance of our contribution, we have trained different simple NN architectures (CNNs, MLPs, and shallow models) on the ESM representation. We kept the best performing model: a three-layer convolutional neural network.

### 3.3 Model Architecture

Similar architectures, like LM-GVP and DeepFRI, are using LM representation at the node/amino acid-level. Whilst this makes intuitive sense, we realized that it leads to scalability issues. Given that the average protein length is 556.90 for the dataset, using one LM representation per amino acid can quickly exceed the average memory of a single GPU or lead to excessive training times. Instead, we are working with a fixed-size representation vector per protein, using the mean to aggregate the representations as suggested by the ESM team.

Our first hypothesis is that the representations from LM and GVP are complementary. Therefore, our first approach was to train two separate classifiers and ensemble them by computing the mean of the output probabilities. The resulting F1-scores were 0.55 for the LM model, 0.46 for the GVP model, and 0.58 for the ensemble. Whilst this ensembling approach already outperformed each classifier separately, we realized that we could improve performance if we trained both classifiers together with a differentiable mean. We believe that this method allows the GVP layers to “learn away” from what the LM already represents. After experimenting with various architectures to combine GVP and LM representations, we found the best performing to be a dense-residual type head (Figure 1.c).

**Figure 1:**
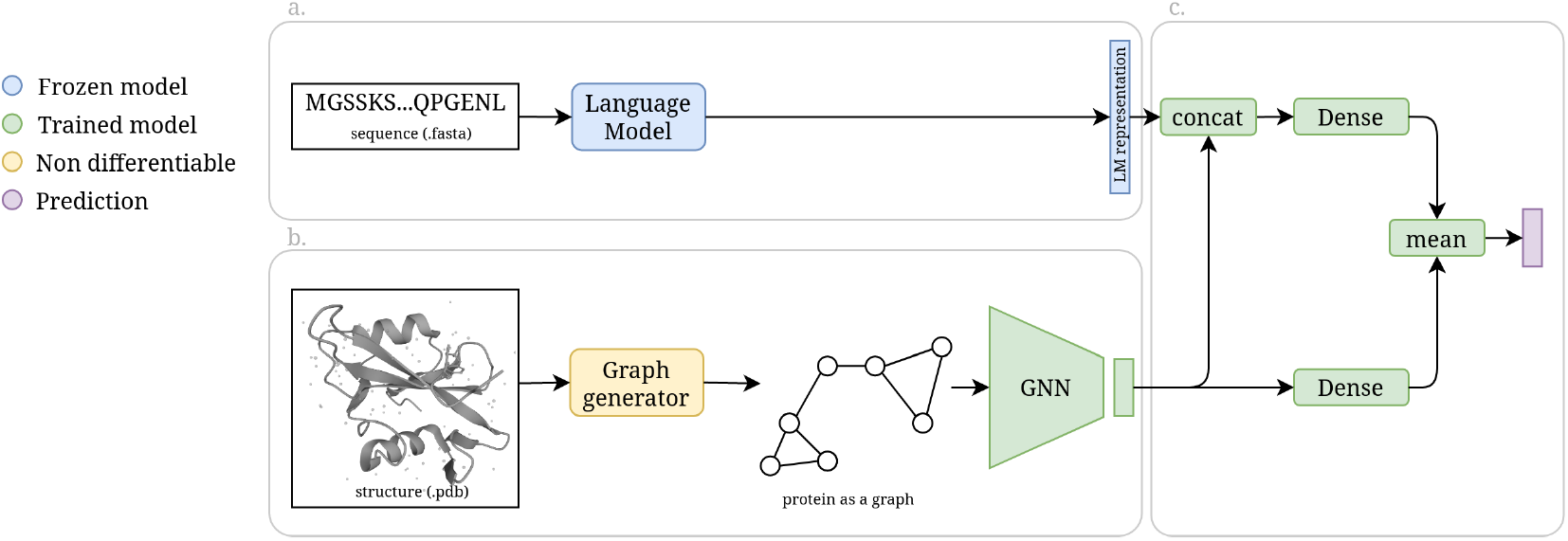
First (a.), a protein sequence is passed to an LM, which outputs a protein representation. Backpropagation does not affect this part as the weights are frozen. In parallel (b.), a PDB structure is processed to generate a graph where nodes represent amino acids and edges are drawn between adjacent amino acids. This graph is used as an input for a GNN (with GVP layers [15]) which outputs a graph representation. Finally (c.), both LM and Graph representations are ensembled through a novel architecture that yields the class probabilities.

## 4 Results

Similar to other researchers[16], we observed that LM-powered models clearly outperform GVP alone. However, our proposed ensembling architecture outperforms the best LM and structure-aware approaches respectively. Note that the proposed architecture (4) using representations at the protein level outperforms the competing model (3) that use representations at the amino acid level; thus provides both faster training and better performance.

**Table 2:** Micro F1-score per class for both LM baseline and our proposed ensembling architecture (Figure 1) on the HPA test set.

| Model | Cytoplasm | Nucleus | Cell memb. | Mitochondrion | Endo. ret. | Golgi |
| --- | --- | --- | --- | --- | --- | --- |
| LM baseline | 0.53 | 0.66 | 0.33 | 0.62 | 0.13 | 0.19 |
| Ours | 0.56 | 0.75 | 0.43 | 0.51 | 0.20 | 0.31 |

We observe that in every class (except Mitochondrion), our proposed architecture outperforms the LM baseline.

In order to show the impact of adding positional encoding as a node feature, we performed an ablation experiment. We trained with and without PE for 60 epochs. Without PE, the test F1-score reached 0.548, with PE it reached 0.569^3^. This validates the hypothesis previously formulated: helping the model extract patterns at specific positions increased the classification performance.

## 5 Discussion

- Given that for mitochondrion, the LM baseline outperforms the proposed architecture, we propose the following explanation. Shuttling proteins into mitochondria is unique: Mitochondrial entry requires a protein to bind to a chaperone while it is in an unfolded form. Upon entry to this organelle, the protein will adopt a folded state. [17]. Hence, we could explain the lower accuracy of GVM by the known dependence on sequence rather than structure features for mitochondrial localization.
- We observe that the prediction of nuclear localization by our ensembling method outperforms DeepLoc’s prediction. As mentioned above, Nuclear localization is frequently mediated by an NLS. However, other routes for nuclear entry exist: a protein without an NLS may interact with other proteins that include an NLS, or other chaperones that mediate entry. These cases are notably more challenging to predict, as they are not mediated directly by a conserved sequence motif such as the NLS. To demonstrate the benefit of incorporating structure information for localization prediction, we focused on cases in which the LM baseline on its own fails in nuclear prediction, but our ensembling method succeeds. We further distilled the list of these cases to include only the cases where DeepLoc provided inaccurate predictions. None of these proteins included an NLS (According to SwissProt annotation [18]), and some were literature-documented cases of proteins that enter the nucleus via non-NLS routes, like General transcription factor IIH subunit 1 (UniProt accession P32780), which is a component of RNA polymerase II. This protein interacts with other proteins to form a complex before entering the nucleus. The complex nuclear entry is then mediated by an NLS motif of other components of the complex [19]. This stands as a promising finding in favor of using structure-aware protein representations.

## 6 Conclusion

In this research, we successfully combined signals from protein structures with Language Model representations to outperform the state of the art of subcellular localization prediction. The principle takeaways are the following: predicted structures appear not to be the sole answer to better protein representations; however, once combined with LM representations and fed into a specific model architecture, such as the one we propose, the quality of representations as well as the scalability of the model improves. Contrasting competing models, we focused on making a scalable/lightweight architecture allowing faster training for more downstream tasks on consumer GPUs, in an effort to democratize structure-aware protein representations models, whilst increasing the model performance.

## Footnotes

1 ProtT5XL has 3B parameters and ESM-1b has 650M parameters. ProtT5XL was shown to be marginally better than ESM-1b for localization prediction. [2]

2 The classes *extracellular* & *plastid* are not present in the test set. Furthermore, we also removed *lysosome/vacuole* & *peroxisome* from the test results since they were not significantly represented with less than 7 occurrences.

3 Furthermore, we observed a better performance during the whole duration of training.

